# Gene copy normalization of the 16S rRNA gene cannot outweigh the methodological biases of sequencing

**DOI:** 10.1101/813477

**Authors:** Robert Starke, Daniel Morais

## Abstract

The 16S rRNA gene is the golden standard target of sequencing to uncover the composition of bacterial communities but the presence of multiple copies of the gene makes gene copy normalization (GCN) inevitable. Even though GCN resulted in abundances closer to the metagenome, it should be validated by communities with known composition as both amplicon and shotgun sequencing are prone to methodological biases. Here we compared the composition of three mock communities to the composition derived from 16S sequencing without and with GCN. In all of them, the 16S composition was different from the mock community and GCN improved the picture only in the community with the lowest Shannon diversity. Albeit with low abundance, half of the identified genera were not present in the mock communities. Our approach provides empirical evidence to the methodological biases introduced by sequencing that was only counteracted by GCN in the case of low α-diversity, potentially due to the small number of bacterial taxa with known gene copy numbers. We thus cannot recommend the use of GCN moving forward and it is questionable whether a complete catalogue of 16S rRNA copy numbers can outweigh the methodological biases of sequencing.

---

Amplicon sequencing of 16S rRNA is the golden standard to describe the composition of bacterial communities due to (i) cost, (ii) availability, (iii) presence of extraction and preparation kits, (iv) taxonomic resolution as deep as the level of genera and (v) previous research. Unsurprisingly it outcompeted (46,473 papers as of February 2019) other possible techniques to describe community structures such as metagenomics (7,699), metaproteomics (367) or metatranscriptomics (439). The general practice as shown by the myriads of publications does not comprise the correction of the obtained raw counts by 16S rRNA gene copy numbers per bacterial genome even though it is known that bacteria can have multiple copy numbers of the gene ^1^. Logically, two bacteria with similar raw counts but different gene copy numbers cannot be equally abundant which is why GCN seems necessary. The recent recommendation against GCN based on the systematic evaluation of the predictability of 16S GCNs in bacteria ^2^ contradict the previous suggestion in favor of GCN based on the comparison of 16S and metagenomics ^1^. However, sequencing techniques are prone to similar methodological biases introduced by extraction, PCR, sequencing and bioinformatics, and could thus similarly diverge from the real picture as recently demonstrated ^3^. We therefore believe that the use of mock communities as standard is inevitable to prove the viability of GCN. Here we compared three randomly chosen taxonomically different mock communities (Mock-2, Mock-20 and Mock-21) from *mockrobiata* provided elsewhere ^4^ that derived from the combination of extracted genomic DNA from bacterial strains and the subsequent 16S rRNA gene amplicon sequencing to estimate the impact of GCN on the bacterial community composition.

Operational taxonomic units (OTU) were annotated by *blastn* ^5^ as best hit down to the genus level with an average similarity match of 97.7±1.7% for Mock-2, 97.4±1.7% for Mock-20 and 97.6±1.6% for Mock-21. In total 3,973 from 34,154 OTU counts (13.2%) in Mock-2 could not be assigned to a bacterial genus compared to 378 from 173,460 (0.22%) for Mock-20 and 328 from 180,542 (0.18%) for Mock-21. Mock-2 comprised of 23 bacterial genera of which only 14 were identified by 16S sequencing opposed to 17 in Mock-20 and 16 in Mock-21 of which all have been identified (**Figure 1**). These findings illustrated missed identifications that seem to be related to the sequencing depth. 30,000 OTU counts for Mock-2 were not sufficient to identify all of the 23 genera in the community whereas 180,000 OTU counts for Mock-20 and Mock-21 resulted in the identification of all genera. However, the three mock communities are simple compared to the billions of organisms belonging to thousands of different species in one gram of soil ^6^. Particularly considering the prokaryotic density of 10,000,000 organisms per gram of soil ^7^ that is at least one magnitude of order higher than per milliliter of water in the ocean ^8^, we conclude that 10,000 but at least 2,000 OTU counts per taxonomic rank of interest are necessary to fully cover the members of the community.

**Figure 1:**
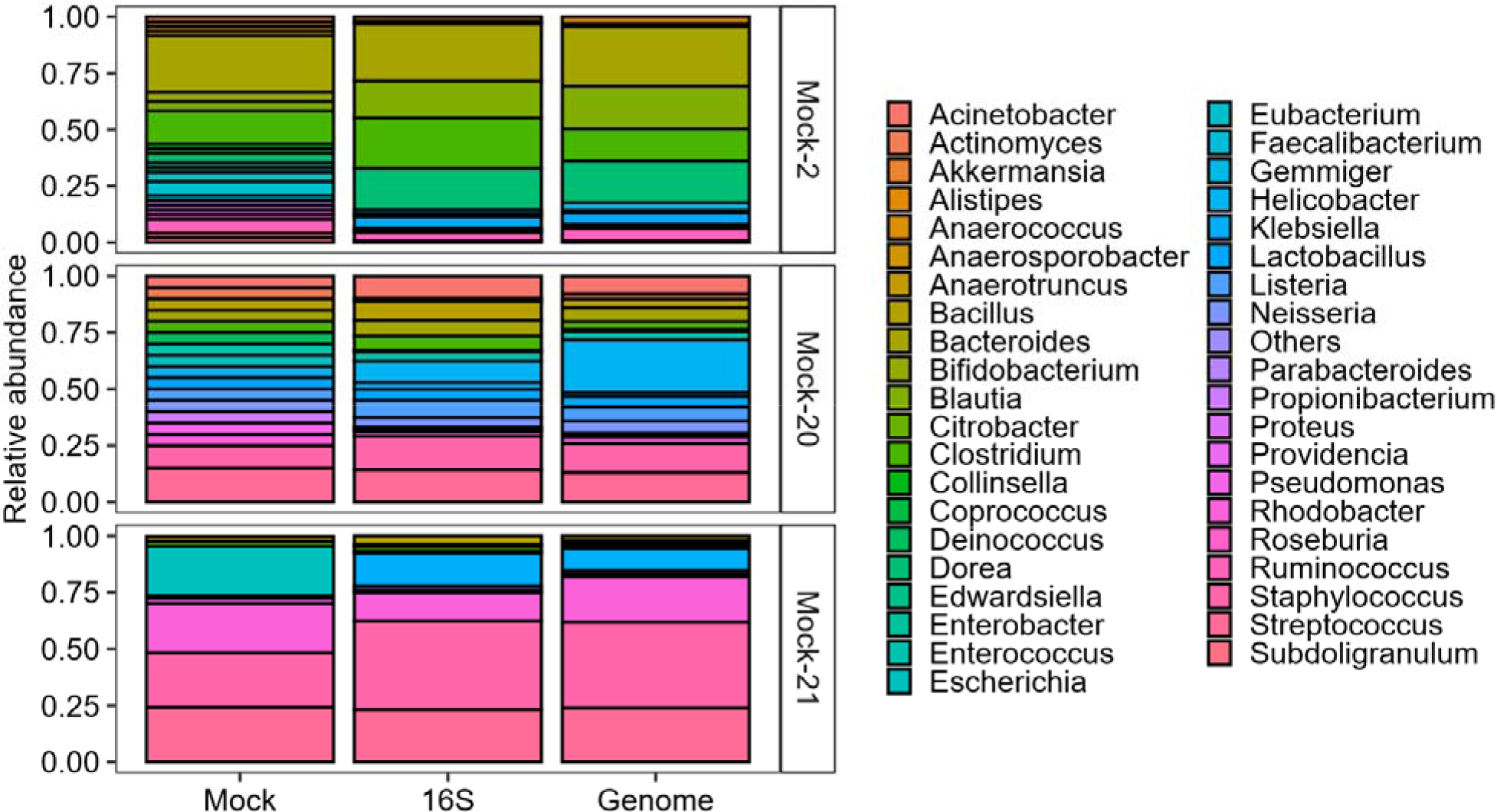
Community structure on the genus level of three mock communities (Mock) and estimated by 16S rRNA sequencing without (16S) and with GCN (Genome).

In total, 19 genera in Mock-2 together with 18 in Mock-20 and 77 in Mock-22 were wrongly identified during sequencing due to their absence when the extracted genomic DNA was combined. Admittedly, the majority among them were found with low abundance, presumably as noise during sequencing. However, *Klebsiella* of the family *Enterobacteriaceae* comprised high abundances in each community. Together with highly abundant extracted DNA from *Escherichia* but low sequencing abundances, we conclude the misidentification of *Escherichia*, also an *Enterobacteriaceae*, as *Klebsiella*. In compliance with our results, phylogenetic trees based on the 16S rRNA gene are ambiguous in *Enterobacteriaceae* and differ in the relative position of several genera ^9,10^. Processes of recombination and gene conversion ^11,12^, and different sequences of the 16S rRNA gene found within a single species ^13^ were previously hold accountable. Here we provide empirical evidence for the misidentification of *Escherichia* as *Klebsiella*, which could prove to be detrimental for proper prophylactic medical treatment since both are pathogens causing a different array of diseases ^14,15^. Even though one advantage of targeting the 16S rRNA gene is taxonomic resolution, we report the failure of correct identification of the genus within *Enterobacteriaceae*, which could be true for other bacterial families as well.

Non-metric multidimensional scaling (NMDS) revealed that the mock community composition derived from combining extracted DNA was indeed different to the composition derived from 16S rRNA gene amplicon sequencing (**Figure 2**). GCN did not impact the community composition and its relative distance within two dimension of the NMDS to Mock-2 and Mock-20, which was further supported by the residual sum of squares between each mock community and 16S sequencing without and with GCN (**Table 1**). However, in Mock-21, GCN resulted in a picture closer to the mock community, presumably due to the low complexity of the community as its Shannon diversity on genus level was 40% lower (1.69) than in Mock-2 (2.71) and in Mock-21 (2.76). Logically we suggest that GCN in communities of low Shannon diversity (≪2.7) could be beneficial. However, the Shannon diversity of bacterial communities typically range between 3 and 6 in both terrestrial ^16–19^ and aquatic ecosystems ^20,21^, which are likely too diverse for GCN to have an impact as shown for Mock-2 and Mock-20.

**Figure 2:**
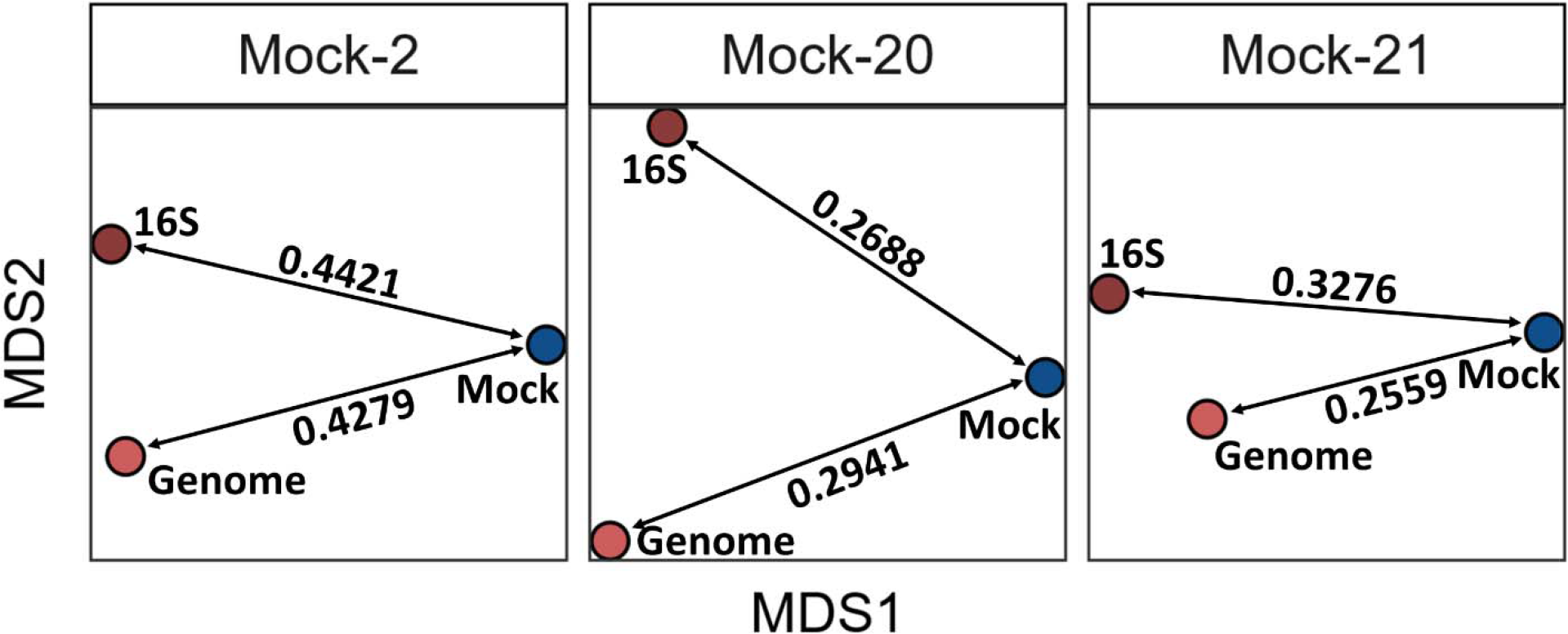
Non-metric multidimensional scaling (NMDS) using Bray Curtis distances in two dimensions of the known community structure (in blue) and derived from 16S rRNA sequencing without (16S) and with GCN (Genome).

**Table 1:** Residual sum of squares (RSS) as discrepancy between the known composition of the mock community and the 16S rRNA sequencing without (16S) and with GCN (Genome) as well as the α-diversity as Shannon diversity on genus level.

| Community | 16S | Genome | α-diversity |
| --- | --- | --- | --- |
| Mock-2 | 0.0576 | 0.0593 | 2.71 |
| Mock-20 | 0.0204 | 0.0466 | 2.76 |
| Mock-21 | 0.0968 | 0.0745 | 1.69 |

Concluding, together with the issues to predict 16S GCNs in bacteria ^2^, we cannot recommend the use of GCN based on the *in vitro* comparison of sequenced amplicons from three randomly chosen mock communities.

## Methods

### Data generation

The community data was obtained from the *mockrobiata* database provided by Bokulich and colleagues ^4^. Three mock communities that contain the reverse reads of sequencing or a clear summary of the known community composition were randomly chosen: Mock-2 that has been described elsewhere ^22,23^ together with Mock-20 and Mock-21 that was obtained through BEI Resources, NIAID, NIH as part of the Human Microbiome Project: Genomic DNA from Microbial Mock Community B (Even, High Concentration), v5.1H, for Whole Genome Shotgun Sequencing, HM-276D. The mock composition generated by the combination of extracted genomic DNA from bacterial strains was downloaded from https://github.com/caporaso-lab/mockrobiota/tree/master/data (Mock-2, Mock-20 and Mock-21) as expected taxonomy using *SILVA* at a 99% identity criterion to remove highly identical sequences ^24^. The raw sequencing data including the forward and reverse reads was processed with *SEED 2* ^25^. Briefly, quality filtering at a cut-off of 30 was followed by clustering of the representative sequences from the clusters as consensus and most abundant, and identification of OTUs by *blastn* ^5^ against a 16S database including chloroplasts and archaea from the ribosomal database project (RDP) as of December 2017 ^26^.

### Gene copy number normalization and statistical analysis

Known 16S rRNA gene copy numbers from bacterial genomes were obtained from the Ribosomal RNA Database (*rrnDB*) as of September 2018 ^27^. Of the 152 genera identified by 16S sequencing, 116 were annotated with a gene copy number on genus level ranging from one to 21 copies from one to 621 genomes per genus with an average gene copy number of 5.54±0.99. For the remaining 36 genera, the next higher taxonomic rank with a gene copy number derived from at was used. For each OTU, The raw counts were divided by the mean gene copy number of the annotated genus to obtain the absolute normalized OTU content. The absolute OTU counts were divided by the total OTU counts to give relative abundances. Non-metric multidimensional scaling (NMDS) using Bray Curtis distances in two dimensions of the known community structure (Mock) and derived from 16S sequencing without (16S) and with GCN (Genome) was performed in *R* using the package *MASS* ^28^. The distance (*d*) between the mock community and the 16S sequencing without (16S) and with GCN (Genome) in two dimensions was derived as straight line between two points (x_1_,y_1_) and (x_1_,y_1_) in a 2D-plane as given by the Pythagorean Theorem (Equation 1). The residual sum of squares (*RSS*) was estimated from the difference of the *i*^*th*^ value between the mock community as *y*_*i*_ and the 16S rRNA sequencing without (16S) and with GCN (Genome) both as *f*(*x*_*i*_) given by Equation 2. The Shannon diversity was calculated on the level of bacterial genera. Visualization was carried out in *R* using the package *ggplot2* ^29^.

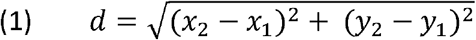

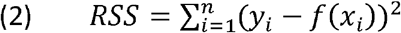

## Acknowledgements

RS thanks the Czech Science Foundation for the project 18-25706S.

## Author contributions

RS and DM designed the experiment. RS and DM performed data analysis. The paper was written by RS and DM. All authors approved the final version of the manuscript.

## References

1. Větrovský, T. & Baldrian, P. The Variability of the 16S rRNA Gene in Bacterial Genomes and Its Consequences for Bacterial Community Analyses. PLoS One (2013). doi:10.1371/journal.pone.0057923

2. Louca, S., Doebeli, M. & Parfrey, L. W. Correcting for 16S rRNA gene copy numbers in microbiome surveys remains an unsolved problem. Microbiome (2018). doi:10.1186/s40168-018-0420-9

3. McLaren, M. R., Willis, A. D. & Callahan, B. J. Consistent and correctable bias in metagenomic sequencing experiments. bioRxiv (2019). doi:http://dx.doi.org/10.1101/559831doi

4. Bokulich, N. A. et al. mockrobiota: a Public Resource for Microbiome Bioinformatics Benchmarking. mSystems (2016). doi:10.1128/mSystems.00062-16

5. BLAST. BLAST Basic Local Alignment Search Tool. Blast Program Selection Guide (2013). doi:10.1006/jmbi.1990.9999

6. Torsvik, V. & Øvreås, L. Microbial diversity and function in soil: From genes to ecosystems. Curr. Opin. Microbiol. (2002). doi:10.1016/S1369-5274(02)00324-7

7. Raynaud, X. & Nunan, N. Spatial ecology of bacteria at the microscale in soil. PLoS One (2014). doi:10.1371/journal.pone.0087217

8. Whitman, W. B., Coleman, D. C. & Wiebe, W. J. Prokaryotes: The unseen majority. Proc. Natl. Acad. Sci. (1998). doi:10.1073/pnas.95.12.6578

9. Granier, S. A. et al. Recognition of two genetic groups in the Klebsiella oxytoca taxon on the basis of chromosomal β-lactamase and housekeeping gene sequences as well as ERIC-1R PCR typing. Int. J. Syst. Evol. Microbiol. (2003). doi:10.1099/ijs.0.02408-0

10. Hedegaard, J., Sperling-Petersen, H. U., Norskov-Lauritsen, N., Steffensen, S. A. d. A. & Mortensen, K. K. Identification of Enterobacteriaceae by partial sequencing of the gene encoding translation initiation factor 2. Int. J. Syst. Bacteriol. (2009). doi:10.1099/00207713-49-4-1531

11. Martinez-Murcia, A. J., Anton, A. I. & Rodriguez-Valera, F. Patterns of sequence variation in two regions of the 16S rRNA multigene family of Escherichia coli. Int. J. Syst. Bacteriol. (2009). doi:10.1099/00207713-49-2-601

12. Hashimoto, J. G., Stevenson, B. S. & Schmidt, T. M. Rates and consequences of recombination between rRNA operons. J. Bacteriol. (2003). doi:10.1128/JB.185.3.966-972.2003

13. Ueda, K., Seki, T., Kudo, T., Yoshida, T. & Kataoka, M. Two distinct mechanisms cause heterogeneity of 16S rRNA. J. Bacteriol. (1999).

14. Chaudhury, A., Nath, G., Tikoo, A. & Sanyal, S. C. Enteropathogenicity and antimicrobial susceptibility of new Escherichia spp. J. Diarrhoeal Dis. Res. (1999).

15. Podschun, R. & Ullmann, U. Klebsiella spp. as nosocomial pathogens: Epidemiology, taxonomy, typing methods, and pathogenicity factors. Clinical Microbiology Reviews (1998).

16. Bastida, F. et al. Differential sensitivity of total and active soil microbial communities to drought and forest management. Glob. Chang. Biol. 23, (2017).

17. Fierer, N. & Jackson, R. B. The diversity and biogeography of soil bacterial communities. Proc. Natl. Acad. Sci. (2006). doi:10.1073/pnas.0507535103

18. Peng, M., Zi, X. & Wang, Q. Bacterial community diversity of oil-contaminated soils assessed by high throughput sequencing of 16s rRNA genes. Int. J. Environ. Res. Public Health (2015). doi:10.3390/ijerph121012002

19. Kaiser, K. et al. Driving forces of soil bacterial community structure, diversity, and function in temperate grasslands and forests. Sci. Rep. (2016). doi:10.1038/srep33696

20. Zhang, H. H. et al. Vertical distribution of bacterial community diversity and water quality during the reservoir thermal stratification. Int. J. Environ. Res. Public Health (2015). doi:10.3390/ijerph120606933

21. Liu, K. et al. Bacterial community changes in a glacial-fed Tibetan lake are correlated with glacial melting. Sci. Total Environ. (2019). doi:10.1016/j.scitotenv.2018.10.104

22. Bokulich, N. A. et al. Quality-filtering vastly improves diversity estimates from Illumina amplicon sequencing. Nat. Methods (2013). doi:10.1038/nmeth.2276

23. Bokulich. A standardlized, extensible framework for optimizing classification improves marker-gene taxonomic assignments. PeerJ Prepr. (2015). doi:10.7287/peerj.preprints.49v1

24. Pruesse, E. et al. SILVA: A comprehensive online resource for quality checked and aligned ribosomal RNA sequence data compatible with ARB. Nucleic Acids Res. (2007). doi:10.1093/nar/gkm864

25. Větrovský, T., Baldrian, P. & Morais, D. SEED 2: A user-friendly platform for amplicon high-throughput sequencing data analyses. in Bioinformatics (2018). doi:10.1093/bioinformatics/bty071

26. Cole, J. R. et al. Ribosomal Database Project: Data and tools for high throughput rRNA analysis. Nucleic Acids Res. (2014). doi:10.1093/nar/gkt1244

27. Stoddard, S. F., Smith, B. J., Hein, R., Roller, B. R. K. & Schmidt, T. M. rrnDB: Improved tools for interpreting rRNA gene abundance in bacteria and archaea and a new foundation for future development. Nucleic Acids Res. (2015). doi:10.1093/nar/gku1201

28. Venables, W. N. & Ripley, B. D. Modern Applied Statistics with S Fourth edition. World (2002). doi:10.2307/2685660

29. Wickham, H. ggplot2: Elegant Graphics for Data Analysis. Journeal Stat. Softw. (2017). doi:10.1007/978-0-387-98141-3

